# The selective 5-HT_2A_ receptor agonist LPH-5 induces persistent and robust antidepressant-like effects in rodents

**DOI:** 10.1101/2024.04.19.590212

**Authors:** Anders A. Jensen, Claudia R. Cecchi, Meghan Hibicke, Astrid H. Bach, Erik Kaadt, Emil Märcher-Rørsted, Charles D. Nichols, Betina Elfving, Jesper L. Kristensen

## Abstract

Psychedelic-assisted psychotherapy has over the last decade emerged as a promising treatment strategy for mental health disease, and the therapeutic potential in classical psychedelics such as psilocybin, LSD and 5-MeO-DMT is presently being pursued in a plethora of clinical trials. However, the resurgent interest in the drugs as therapeutics has also prompted a search for novel agents with more specific pharmacological activities than the rather promiscuous classical psychedelics. Here we present the results of an elaborate preclinical characterization of one such compound, LPH-5 [(*S*)-3-(2,5-dimethoxy-4-(trifluoromethyl)phenyl)piperidine]. LPH-5 was found to be a potent partial agonist at the 5-HT_2A_ receptor (5-HT_2A_R) and to exhibit pronounced selectivity for this receptor over the related 5-HT_2B_ and 5-HT_2C_ receptors in a range of functional assays. LPH-5 (0.375 – 12.0 mg/kg, *i.p.*) dose-dependently induced head-twitch responses (HTR) in Sprague Dawley rats, with substantial 5-HT_2A_R engagement being observed at 0.5-1.0 mg/kg. Acute administration of LPH-5 (1.5 mg/kg, *i.p*.) induced robust antidepressant-like effects in Flinders Sensitive Line rats and adrenocorticotropic hormone-treated Sprague Dawley rats, and LPH-5 (0.3 and 1.5 mg/kg, *i.p*.) induced significant effects in a recently developed Wistar Kyoto rat model proposed to reflect the long-term antidepressant-like effects produced by psychedelics in humans. In conclusion, selective 5-HT_2A_R activation, as mediated here by LPH- 5, seems to hold antidepressant potential, suggesting that this activity component is key for the beneficial effects of classical psychedelics. Hence, we propose that LPH-5 and other 5-HT_2A_R- selective agonists could hold potential as therapeutics in psychiatric disease as a new generation of psychedelic-derived antidepressant.

## INTRODUCTION

Psychedelic-assisted psychotherapy has emerged as a promising treatment strategy for psychiatric disease.^1–7^ The classical psychedelics like psilocybin, lysergic acid diethylamide (LSD), and *5*-methoxy-*N,N*-dimethyltryptamine (5-MeO-DMT) that presently are under clinical development for these indications are rather promiscuous drugs that possess roughly equipotent activity at several serotonergic receptors and in some cases also at other monoaminergic receptors and/or other receptor types.^8–12^ Concomitantly with the clinical development of the classical psychedelics, a substantial amount of research has thus been dedicated to delineate the molecular, cellular and neural basis for their beneficial effects.^13^ These investigations have spawned several hypotheses about the underlying mechanisms of action of the psychedelics and about the importance of specific serotonin receptor subtypes,^14–16^ heterodimeric receptor complexes,^12^ other receptors,^10^ intracellular signaling cascades,^11,17,18^ and intracellular receptors^19^ for these beneficial effects.

The resurgent interest in psychedelics and the various hypotheses for the origin of their effects have also prompted search for drugs with distinct pharmacological profiles compared to the psychedelics. Amongst the receptors targeted by classical psychedelics the 5-HT_2A_ serotonin receptor (5-HT_2A_R) has so far attracted the most attention as a putative therapeutic target. As the major postsynaptic serotonin receptor in the CNS, 5-HT_2A_R plays key roles for cognitive functions, mood, appetite and circadian rhythm.^14,20–22^ While substantial evidence links the psychedelic effects of classical psychedelics to their shared 5-HT_2A_R agonist activity, it is up for discussion whether this activity component also is key for their beneficial effects on psychiatric diseases, given the lack of clinical investigations with selective 5-HT_2A_R agonists.^13^ This has prompted pursuits into whether the beneficial effects of psychedelics could be retained in drugs that do not possess 5-HT_2A_R agonist activity. Conversely, the possibility that the beneficial effects of psychedelics may arise, fully or in part, from their 5-HT_2A_R activation has also spawned the search for G-protein- and β-arrestin-biased agonists for the receptor, but it remains to be unequivocally delineated whether the different behavioral effects induced by psychedelics are indeed attributable to distinct 5-HT_2A_R signaling pathways.^11,17,18^

The high conservation of orthosteric sites in serotonin receptors, in particular in the three 5- HT_2_R subtypes 5-HT_2A_R, 5-HT_2B_R and 5-HT_2C_R, has made it challenging to develop truly selective 5-HT_2A_R agonists.^23^ While only a few selective 5-HT_2A_R agonists have been developed, 25CN-NBOH being the prototypic 5-HT_2A_R-selective agonist,^24,25^ recent high resolution receptor structures have facilitated structure-based design of subtype-selective and/or biased agonists.^18,26,27^ However, as no 5-HT_2A_R-selective agonist has been entered into clinical trials to date, the therapeutic potential in specific augmentation of 5-HT_2A_R signaling and whether the clinical effects arising from this significantly differ from those induced by the classical psychedelics thus remains to be elucidated.^8^

Recently we reported the discovery of a novel class of 2,5-dimethoxyphenylpiperidine-based 5- HT_2_R agonists and characterized their structure-activity relationship at 5-HT_2A_R and 5- HT_2C_R.^28,29^ The analog LPH-5 [(*S*)-3-(2,5-dimethoxy-4-(trifluoromethyl)phenyl)piperidine, Fig. 1A] exhibited a highly interesting functional profile in a Ca^2+^ imaging assay, acting as a potent partial agonist at 5-HT_2A_R (EC_50_: 3.2 nM), while exhibiting 60-fold lower agonist potency at 5- HT_2B_R and so negligible agonist efficacy at 5-HT_2C_R that it was a *de facto* competitive antagonist at this receptor (IC_50_: 320 nM).^28^ Moreover, LPH-5 possessed 10-fold higher binding affinity to 5-HT_2A_R than to 5-HT_2B_R and 5-HT_2C_R in [^125^I]DOI competition binding assays, and it displayed at least 100-fold higher binding affinity to 5-HT_2A_R than to a range of other targets, including 8 other serotonin receptors and various dopaminergic, adrenergic, histaminergic and muscarinic receptors.^28^

**Fig. 1.**
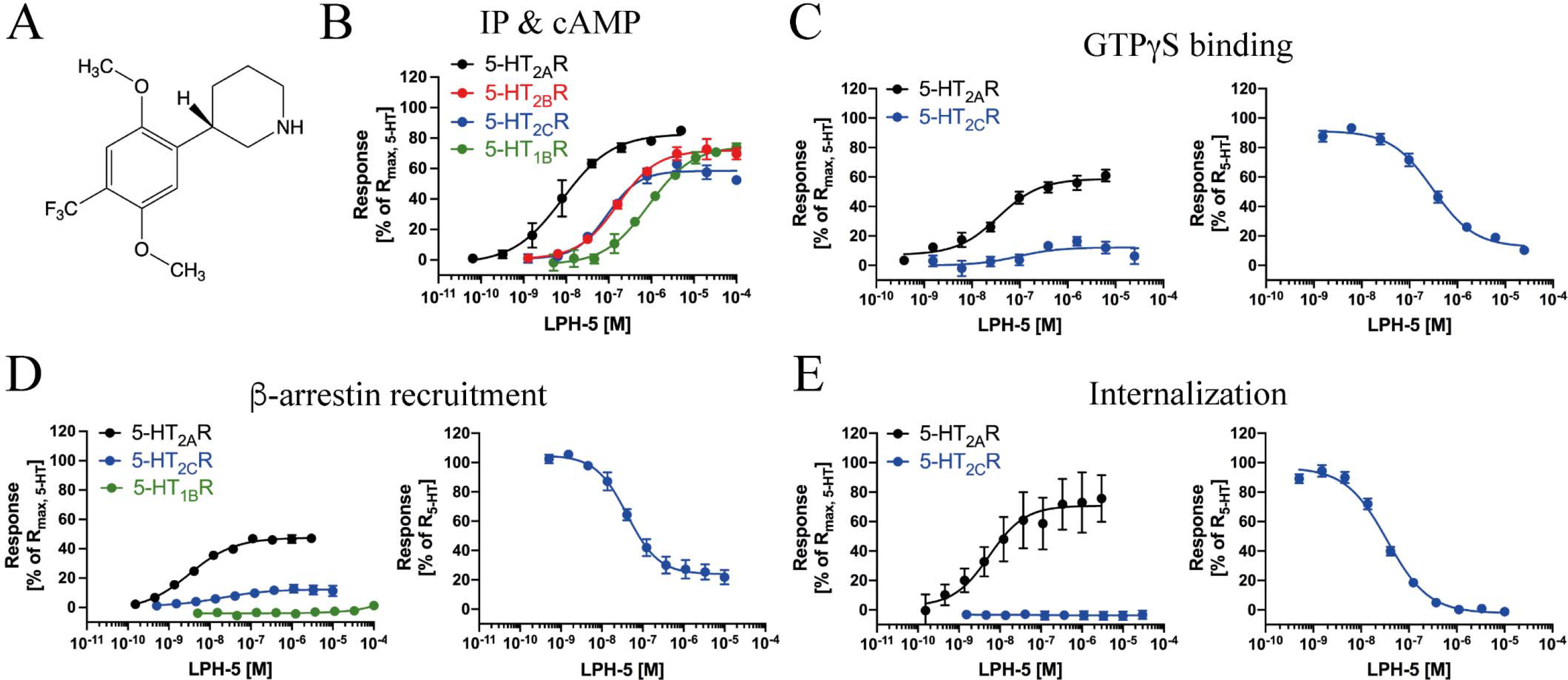
Functional properties displayed by LPH-5 at recombinant human 5-HT_2A_R, 5- HT_2B_R, 5-HT_2C_R and 5-HT_1B_R. (A) Chemical structure of LPH-5. **(B)** Concentration-response relationships displayed by LPH-5 at 5-HT_2A_R, 5-HT_2B_R and 5-HT_2C_R in inositol phosphate (IP) assays and at 5-HT_1B_R in a HitHunter® cAMP assay. **(C)** Concentration-response relationships displayed by LPH-5 at 5-HT_2A_R and 5-HT_2C_R *(left)* and concentration-inhibition relationship displayed by LPH-5 tested in antagonist mode at 5-HT_2C_R (*right)* in GTPγS binding assays. **(D)** Concentration-response relationships displayed by LPH-5 at 5-HT_2A_R, 5-HT_2C_R and 5-HT_1B_R *(left)* and concentration-inhibition relationship displayed by LPH-5 tested in antagonist mode at 5-HT_2C_R (*right)* in PathHunter® β-arrestin recruitment assays. **(E)** Concentration-response relationships displayed by LPH-5 at 5-HT_2A_R and 5-HT_2C_R *(left)* and concentration-inhibition relationship displayed by LPH-5 tested in antagonist mode at 5-HT_2C_R (*right)* in internalization assays. **(B-E)** Data are given as mean ± S.E.M. values based on at least three independent determinations performed in duplicate, except for the 5-HT_1B_R data that are based on independent determinations performed in duplicate. The average pharmacological data extracted from the experiments are given in Table 1.

**Table 1.**
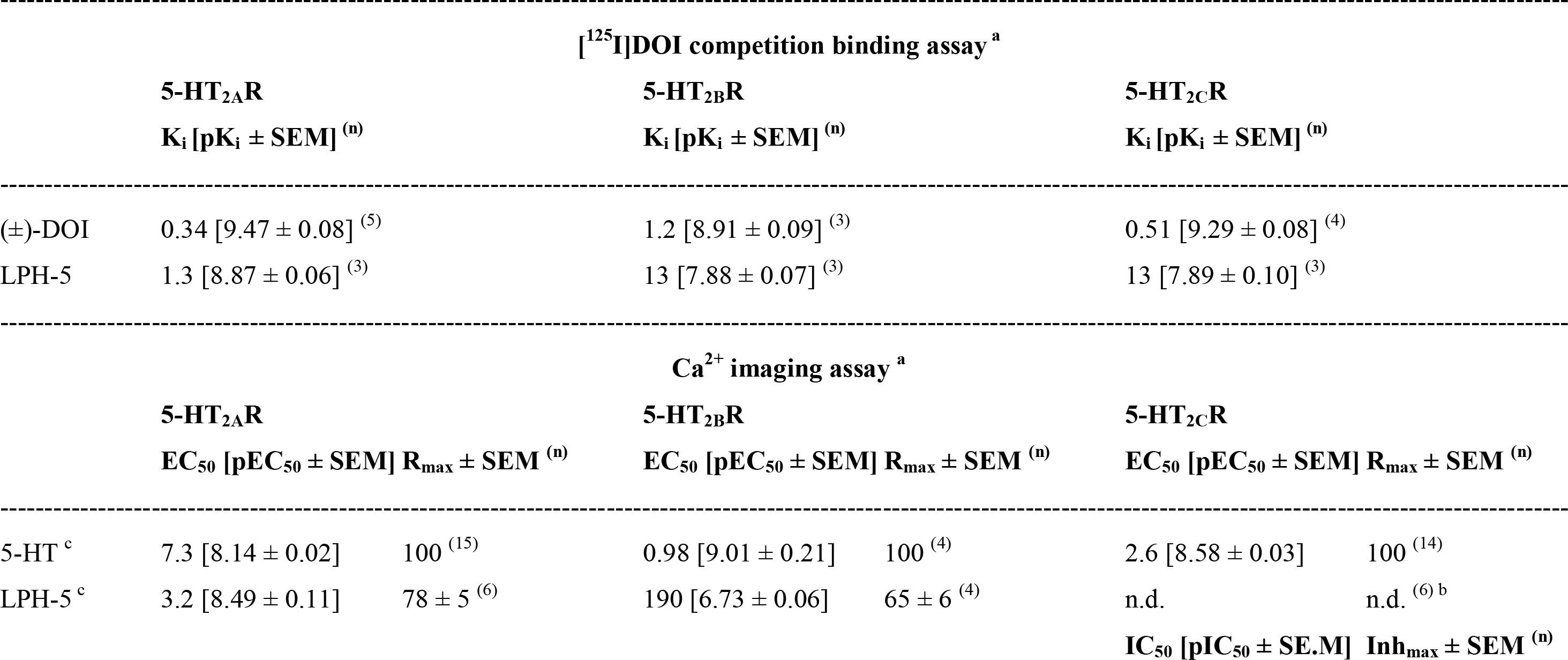
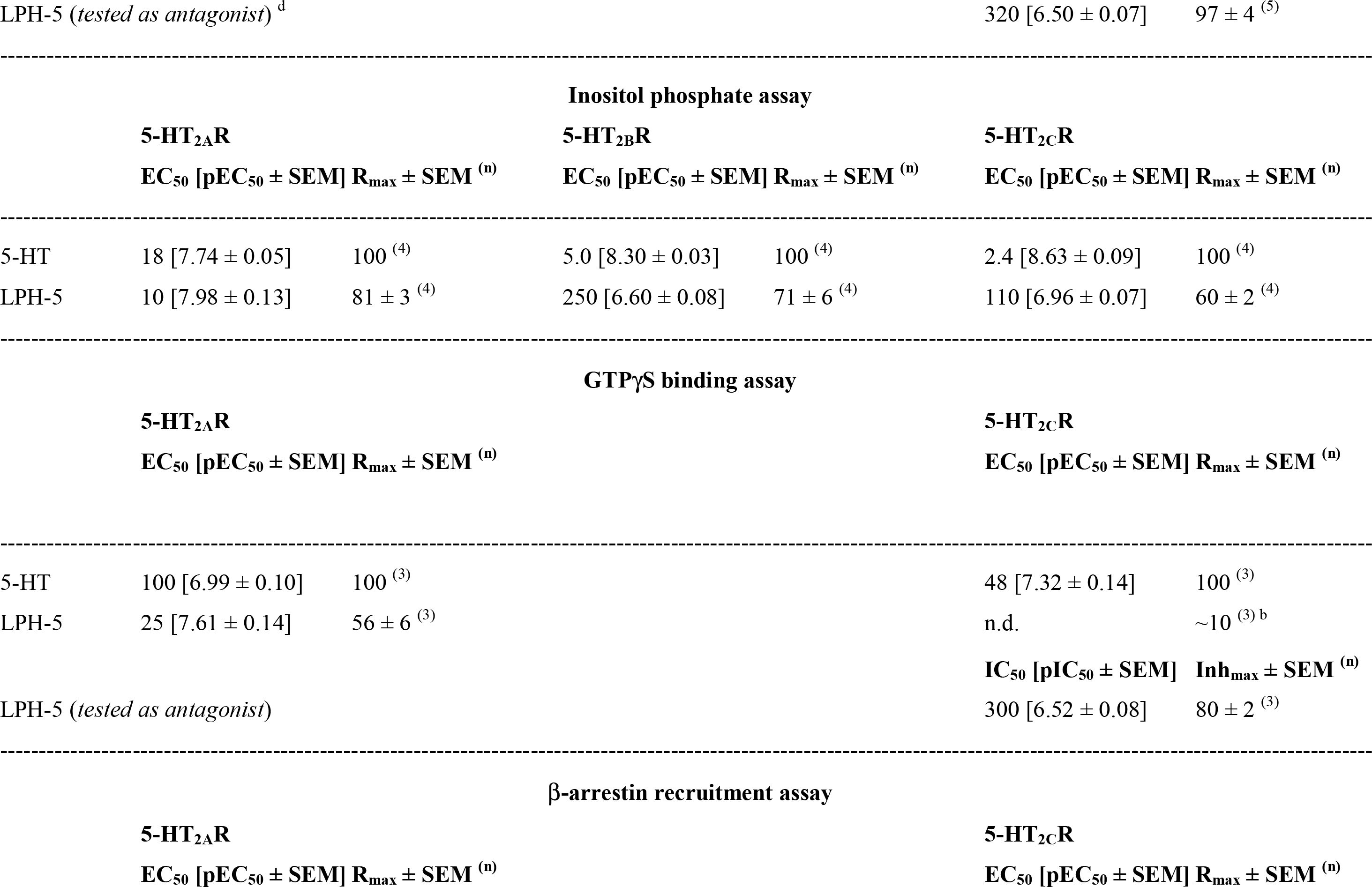
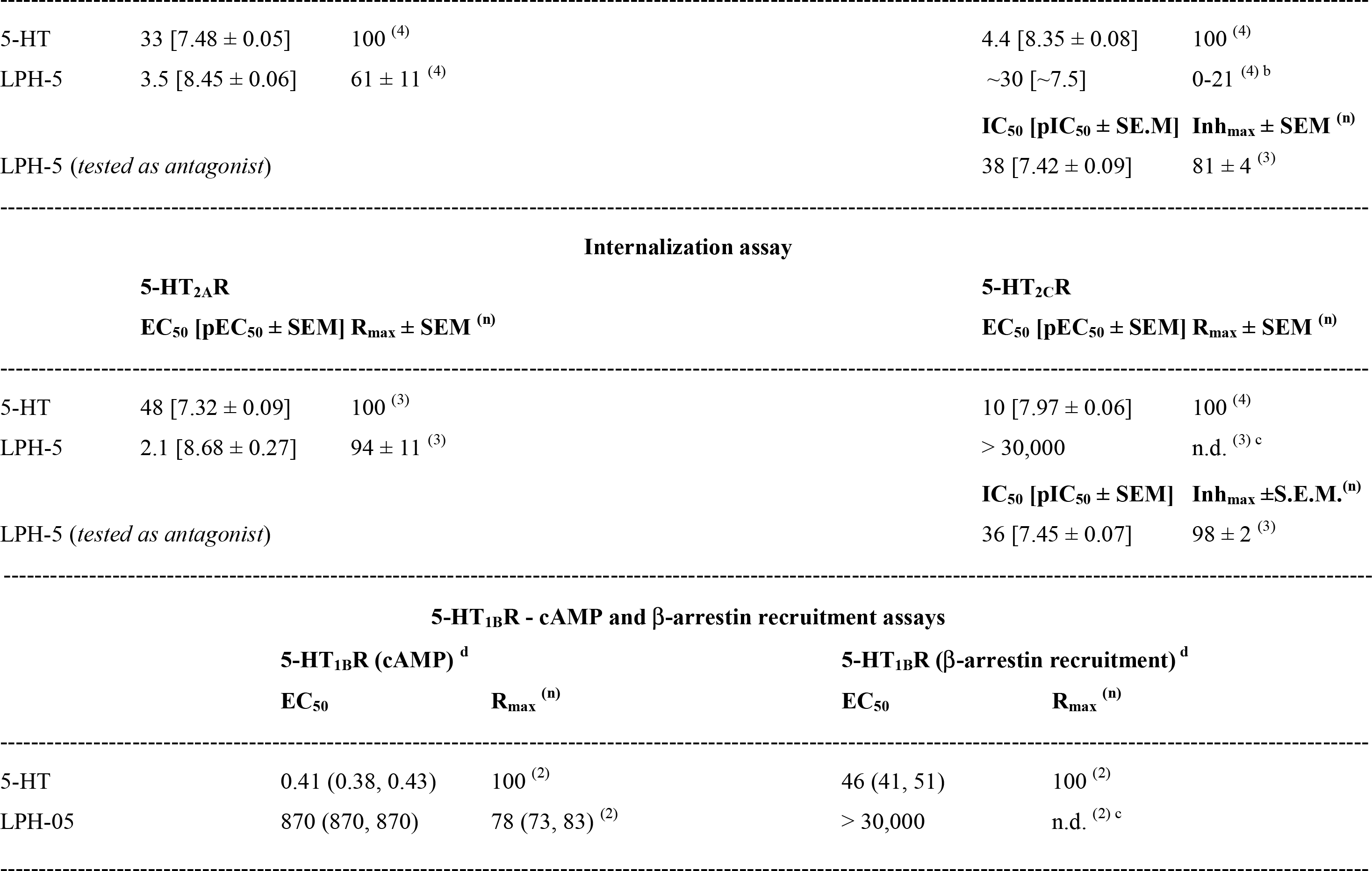
Pharmacological properties exhibited by LPH-5 at the recombinant human 5-HT_2A_R, 5-HT_2B_R, 5-HT_2C_R and 5-HT_1B_R expressed in mammalian cell lines in various binding and functional assays. K_i_ and EC_50_ values are given nM with pK_i_ ± S.E.M. and pEC_50_ ± S.E.M. values in brackets, respectively, and R_max_ (± S.E.M.) values are given as % of 5-HT R_max_ at the receptor. In the functional assays for 5-HT_2C_R, where LPH-5 also was tested in antagonist mode at the 5-HT-induced response through the receptor, IC_50_ values are given nM with pIC_50_ ± S.E.M. values in brackets, and the maximal degree of inhibition (Inh_max_ ± S.E.M.) mediated by LPH-5 at saturating concentrations are given as % of complete inhibition of the 5-HT-induced response. The properties of (±)-DOI (in the binding assay) and 5-HT (in the functional assays) are given for reference. The numbers of independent experiments (n) performed in duplicate underlying the data are given in superscript.

In the present study we have performed a detailed functional profiling of LPH-5 at the human 5- HT_2_Rs, investigated its bioavailability and target occupancy *in vivo* by assessing its induction of head twitch responses in rats, and delineated its therapeutic potential in three rat models of depression.

## MATERIALS AND METHODS

### FUNCTIONAL ASSAYS

#### Assays

The functional properties of LPH-5 at human 5-HT_2A_R, 5-HT_2B_R, 5-HT_2C_R and 5- HT_1B_R were characterized in five different functional assays by Eurofins: in IP-One HTRF® assays,^30,31^ in a fragment complementation-based cAMP assay,^32,33^ in scintillation proximity assay-based GTPγS binding assays,^34,35^ in PathHunter® β-arrestin assays,^36^ and in PathHunter® internalization assays.^37^ The specific assay details are given in detail in Supplemental Information.

Data analysis was performed in GraphPad Prism 10.0 (GraphPad Software, San Diego, CA). Concentration-response and concentration-inhibition curves were fitted to nonlinear regression curves fit with variable slopes: *Y = Bottom + (Top-Bottom)/(1+10^^^((LogEC_50_-X)*n_H_))* and *Y = Bottom + (Top - Bottom)/(1+10^((LogIC_50_-X)*n_H_)*, respectively, where Y is the response, Top and Bottom values are plateaus in the units of the response axis, X is the test compound concentration, EC_50_ is the test compound concentration that yields a response half way between Bottom and Top, IC_50_ is the test compound concentration yielding inhibition to a response half way between Bottom and Top, and n_H_ is the Hill slope.

### ANIMAL EXPERIMENTATION AT AARHUS UNIVERSITY

#### Animals

Male Sprague Dawley (SD) rats were obtained from Taconic (Ry, Denmark). Male Flinders Sensitive Line (FSL) rats and their control Flinders Resistant Line (FRL) rats (derived from the colony at University of North Carolina, NC). The animals were housed in pairs (Cage 1291H Eurostandard Type III H, 425 x 266 x 185 mm, Techniplast, Buguggiate, Italy) at 20 ± 2°C on a 12 h light/dark cycle with ad libitum access to chow pellets and tap water, and the welfare of the animals was assessed daily by animal caretakers. All experimental procedures were performed in specially equipped rooms within the vivarium. All animal procedures were approved by the Danish National Committee for Ethics in Animal Experimentation (2016-15- 0201-01105).

#### Head Twitch Response (HTR) measurements

A total of 56 male SD rats aged 8-9 weeks and weighing 276 ± 22 g (mean ± S.D.) were used. The animals were habituated to the facility for 3 weeks before the experiments. DOI [(±)-2,5-dimethoxy-4-iodoamphetamine] (Sigma Aldrich, St. Louis, MO) and LPH-5^28^ were dissolved in sterile 0.9% saline solution (Fresenius lab, Bad Homburg v.d.H, Germany). The rats were divided into seven experimental groups (n = 8 per group) and injected intraperitoneally (*i.p*.) with vehicle (0.9% saline solution), DOI (3.0 mg/kg), or 5 different doses of LPH-5 (0.375, 1.5, 3.0, 6.0, or 12.0 mg/kg) in a final volume of 1.0 - 1.5 ml/kg.

After drug administration the rats were immediately moved to a standard cage (425 x 266 x 185Lmm) without enrichment material and videotaped for 180 min both from above and from the front. Rats injected with DOI (3.0 mg/kg) or LPH-5 (1.5, 3.0, 6.0, or 12.0 mg/kg) that displayed no or very few head twitches (compared to the other animals in the same experimental group) in the recorded period were classified as outliers and excluded from the final dataset presented. However, the complete HTR data set for all rats is given in Supplemental Information Fig. 1.

#### The Flinders rat model for depression

Male FSL (n=28) and FRL (n=10) rats (9–11 weeks of age; 228–336 g) were used for the experiments. Both *S*-ketamine and LPH-5 were dissolved in sterile 0.9 % saline. The following groups were included in the experiment: FRL-saline (n=10), FSL-saline (n=10), FSL-ketamine (15 mg/kg *S*-ketamine, Pfizer) (n=8), and FSL-LPH-5 (1.5 mg/kg) (n=10).

*Open field test (OFT).* 50 min after injection locomotor activity in the rats was tested in an open field to assess possible inhibitory or stimulatory drug effects that could confound the performance in the forced swim test. A squared open field arena (plastic; 50 × 50 × 37 cm) with a light intensity of approximately 5 lux was used. Each rat was allowed to move freely for 5 min. The test arena was thoroughly cleaned with 70% ethanol between each rat to minimize the impact of olfactory cues. The session was recorded with a camera located directly above the center of the field, and total distance moved was quantified using EthoVision XT video tracking software (version 11.0.928; Noldus Information Technology, Wageningen, The Netherlands).

*Forced swim test (FST).* After the OFT and 60 min after the injection of either saline, *S-* ketamine, or LPH-5, the antidepressant-like effect was assessed in a modified FST.^38^ Since FSL rats inherently show a depression-like phenotype, no pre-swim session was required. The rat was placed in an acrylic plastic cylinder (60 cm in height, 24 cm in diameter) containing 40 cm of water (25±1 °C) for 7 min. A camera located directly in front of the cylinder recorded the session. An experienced investigator blinded to the treatments measured the time spent climbing (defined as attempts to climb the cylinder wall or diving), swimming (defined as a forward propulsion in the water surface) and being immobile (defined as the absence of movements except for the necessary activities to keep the head above water).

#### Adrenocorticotropic hormone (ACTH) model for depression

Male SD rats (age 7 weeks and weight 181-230 g at the beginning of the experiment; age 9 weeks and weight 246-352 g at behavioral testing, n=70) were used in the experiments that were performed essentially as previously described.^39,40^ After 1 week habituation, the rats were administered either 100 µg ACTH/day *s.c.* [ACTH 1-24 (China Peptides, China)] (n=50) or saline (0.9 % saline) (n=20) at 10 a.m. for 14 consecutive days. At day 14, the rats were subjected to a pre-swim test (15 min) after which the rats in the ACTH group (n=50) were administered with either 3 injections of imipramine (n=10) (IMI, 15 mg/kg i.p. (Sigma-Aldrich, Denmark)) or 1 injection of saline, psilocybin (1 mg/kg, i.p. (THC Pharm, Frankfurt, Germany)), psilocybin (10 mg/kg, i.p.), or LPH-5 (1.5 mg/kg, i.p.) (n=10/group). The rats in the saline group (n=20) were administered with either 3 injections of imipramine (n=10) or saline (n=10), thus creating 7 groups with 10 animals in each group (SAL-SAL, SAL-IMI, ACTH-SAL, ACTH-IMI, ACTH-PSI-1, ACTH-PSI-10, ACTH-LPH-5). The 3 imipramine-injections were given at 24 h, 18 h, and 1 h prior to the FST at day 15, whereas psilocybin and LPH-5 were injection 1 h prior to the FST. Preceding the FST, rat baseline mobility was determined in the OFT. The OFT and FST were conducted as described above.

#### Data and Statistical analysis

Results are presented mean ± S.E.M., and significance is defined as p < 0.05. Data and statistical analyses were performed using GraphPad Prism version 10. Outliers were tested for with ROUT-test (significance level Q = 0.01), and no outliers were identified. For the Flinders rat data, one-way ANOVA was used to assess variance across groups followed by a post hoc Dunnett’s multiple comparisons test, comparing FRL-Saline and each treatment group (FSL-ketamine and FSL-LPH-5) to the FSL-Saline group. For the ACTH rat model data, two-way ANOVA followed by Sidak’s multiple comparisons test was used to compare control groups (SAL-SAL, SAL-IMI, ACTH-SAL, ACTH-IMI) to confirm interaction between ACTH treatment and imipramine treatment. One-way ANOVA followed by Dunnett’s multiple comparisons test was used to compare treatment effects for ACTH-treated animals (ACTH-SAL, ACTH-IMI, ACTH-PSI-1, ACTH-PSI-10, and ACTH-LPH-5).

### ANIMAL EXPERIMENTATION AT LOUISIANA STATE UNIVERSITY

#### Animals

Male Wistar Kyoto rats (6 weeks old at start) were obtained from Charles River Laboratories and allowed to habituate to the colony room for at least two weeks prior to experimentation. The animals were housed in pairs in standard modified barrier cages with wood chip bedding on a 12/12 light/dark cycle; ∼70% humidity and 72°F, ad libitum access to water and standard rodent chow. All animal care and experimental procedures were approved by the Institutional Animal Care and Use Committee at Louisiana State University Health Sciences Center.

#### Drugs and study design

All drugs were dissolved in sterile physiological saline at a concentration of 1 mg/ml. Drug was administered *i.p.* at a volume of 1 ml/kg, and each group [saline; 1.0 mg/kg psilocybin (Cayman Chemical, Ann Arbor, MI); 0.3 mg/kg LPH-5; 1.5 mg/kg LPH-5] comprised 8 animals. Following drug treatment, animals were maintained in their home cage for 35 days prior to the behavioral testing.

##### OFT

Locomotor activity (LCA) was assessed in an open field arena immediately prior to each FST to account for any potential confounding by nonspecific sedative or stimulant effects. LCA assessment was performed by placing each rat into a square open field arena (61 cm^2^) with opaque walls 45 cm high and allowing the rats to explore freely for 5 min. The rats’ movements within the arena were recorded and later scored for distance travelled (cm) using EthoVision XT 8.5 tracking software (Noldus Information Technology).

##### FST

Rats were placed into a cylindrical plastic tank (114 cm x 30.5 cm) that contained 30 cm water at 28-30°C. Fresh water was used for each animal. The water depth was such that the rats could not support themselves by touching the bottom of the tank with their hind paws, and their tails could not touch the bottom of the tank while keeping their noses above water. A five-minute swim was video recorded and later scored for immobility, swimming, and climbing by an observer blinded to the treatment. Immobility was defined as no active attempts to escape while maintaining a floating posture in which the rats make only the movements necessary to keep their heads above water. Swimming was defined as actively attempting escape with motions directed outward against the wall of the cylinder. Climbing was defined as actively attempting escape with motions directed upward against the wall of the cylinder.

#### Statistical analysis

The data generated were analyzed using one-way ANOVA with Holm-Sidak post hoc.

## RESULTS

### LPH-5 is a potent and highly selective 5-HT_2A_R agonist

In this work, our preliminary pharmacological characterization of LPH-5 in a recent study^28^ was elaborated with functional profiling of the drug at 5-HT_2A_R, 5-HT_2B_R and 5-HT_2C_R in four other functional assays (Fig. 1B-E, Table 1). The functional properties of LPH-5 were also characterized at 5-HT_1B_R, as a representative of the “non-5-HT_2_R” serotonin receptors.

In an inositol phosphate assay, LPH-5 mediated robust partial agonism at all three 5-HT_2_Rs, displaying 25- and 11-fold lower agonist potencies at 5-HT_2B_R and 5-HT_2C_R than at 5-HT_2A_R, respectively. In a cAMP assay, an analogous second messenger assay, LPH-5 displayed 87-fold lower agonist potency at 5-HT_1B_R than that at 5-HT_2A_R in the inositol phosphate assay (Fig. 1B, Table 1). LPH-5 was also a potent partial agonist at 5-HT_2A_R in the GTPγS binding assay, whereas it displayed significant but so low agonist efficacy at 5-HT_2C_R that a reliable EC_50_ could not be determined for it (Fig. 1C *left*). When tested as an antagonist at 5-HT_2C_R in the GTPγS binding assay, LPH-5 inhibited the 5-HT-induced receptor response in a concentration-dependent manner displaying a high-nanomolar IC_50_ value, but in concordance with it with being a low-efficacious agonist LPH-5 did not eliminate the 5-HT-induced response completely (Fig. 1C *right*, Table 1). This profile was mirrored in a β-arrestin recruitment assay, where LPH-5 was a potent partial 5-HT_2A_R agonist (EC_50_ 3.5 nM, R_max_ 61%) but exhibited a so low R_max_ value at 5-HT_2C_R that its potency only could be roughly estimated (EC_50_ ∼30 nM) (Fig. 1D *left*). When tested in antagonist mode at 5-HT_2C_R in the β-arrestin recruitment assay, LPH-5 inhibited the 5- HT-induced response through the receptor in a concentration-dependent manner (Fig. 1D *right*). Interestingly, LPH-5 also displayed negligible agonist efficacy at 5-HT_1B_R in the β-arrestin recruitment assay (Fig. 1D *left*). Finally, LPH-5 was found to be an efficacious agonist when it came to its induction of 5-HT_2A_R internalization, whereas it did not induce significant 5-HT_2C_R internalization at concentrations up to 30 μM (Fig. 1E *left*). LPH-5 inhibited the 5-HT-induced internalization of 5-HT_2C_R, with saturating concentrations of the compound eliminating the receptor internalization completely (Fig. 1E *right*).

### LPH-5 induces robust head twitch response in rats

HTR, a rhythmic paroxysmal side-to-side head movement in rodents, is a behavioral signature of serotonergic psychedelics.^41,42^ Taking advantage of the correlation between 5-HT_2A_R activation and this behavior, we used the HTR induced by LPH-5 as a read-out of *in vivo* 5-HT_2A_R engagement to assess its dose-response relationship and pharmacokinetic properties in rats. The HTR produced by saline, five LPH-5 doses, and a high dose of the reference 5-HT_2_R agonist DOI in male SD rats determined in 10-min intervals over a 180 min period following *i.p.* administration are presented in Fig. 2.

**Figure 2.**
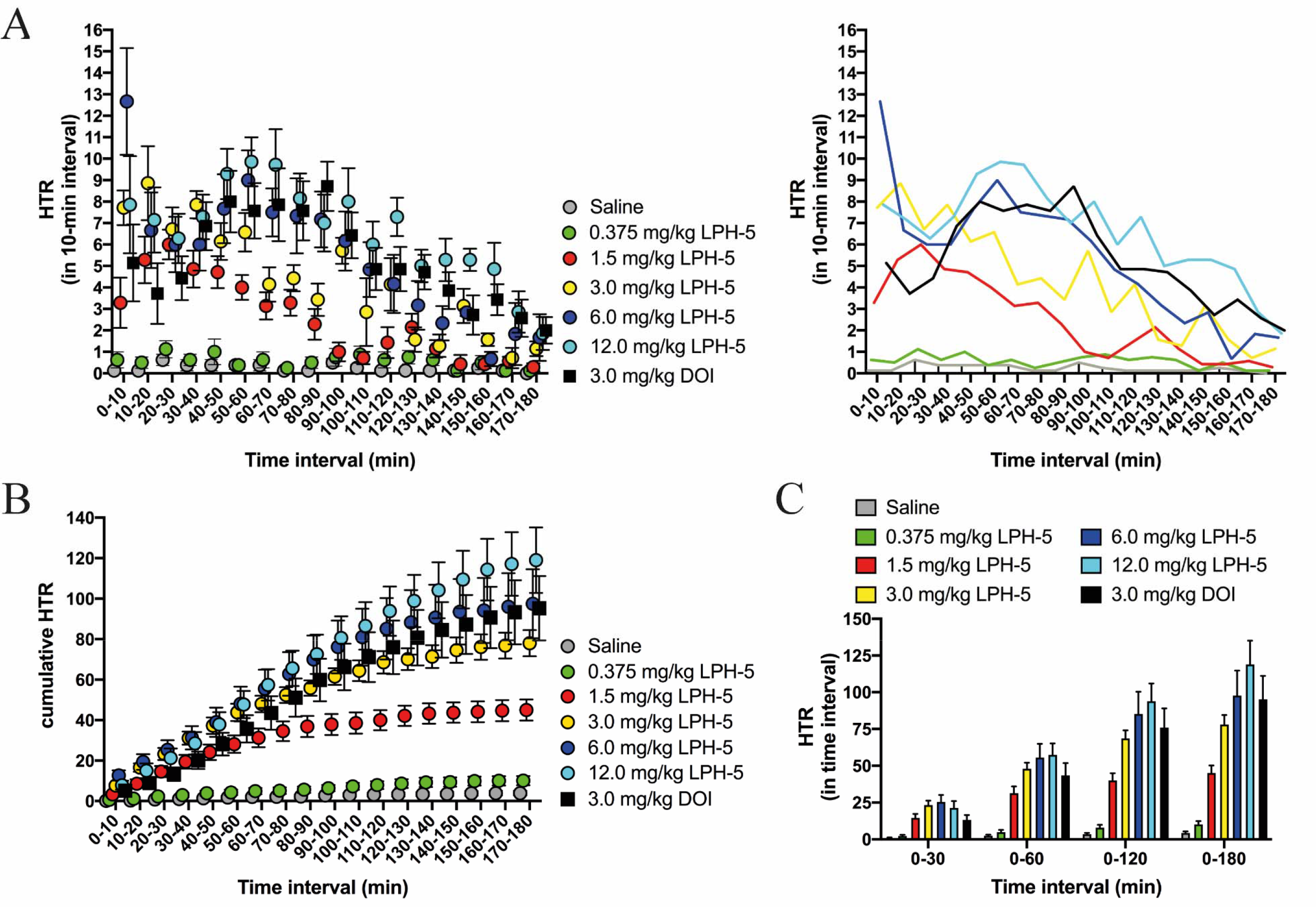
Head twitch response (HTR) induced by *i.p.* injections of saline, LPH-5 (0.375; 1.5; 3.0; 6.0; 12.0 mg/kg) and DOI (3.0 mg/kg) in male Sprague Dawley rats. (A) Time course of the HTR induction in the rats. Average HTR numbers recorded in the rats for each 10- min interval over a total period of 180 min following the injection are given as mean ± S.E.M. values (*left*) and as curves (*right*). (**B)** Cumulative HTR induced in the rats. Cumulative HTR numbers recorded in the rats over a total period of 180 min following the injection are given as mean ± S.E.M. values. **(C)** Dose-effect relationship for the induced HTR in the rats. Average numbers of HTR recorded in the rats in four different time intervals (0-30, 0-60, 0-120, 0-180 min) are given as mean ± S.E.M. values. **(A-C)** The data are based on n=8 [saline and 0.375 mg/kg LPH-5], n=7 [1.5 mg/kg LPH-5; 3.0 mg/kg LPH-5] and n=6 [6.0 mg/kg LPH-5; 12.0 mg/kg LPH-5] animals. The HTR measurements in a total of six rats (one dosed with 1.5 mg/kg LPH-5; one dosed with 3.0 mg/kg LPH-5; two dosed with 6.0 mg/kg LPH-5; two dosed with 12.0 mg/kg LPH-5) were deemed to constitute outlier data and are not included in this figure. However, the HTR raw data for all 56 animals, including the six outlier animals, are given in Supporting Information Fig. 1.

Whereas the lowest LPH-5 dose (0.375 mg/kg) mediated very modest increases in HTR counts compared to the saline control, the four other LPH-5 doses (1.5; 3.0; 6.0; 12.0 mg/kg) and DOI (3.0 mg/kg) all evoked robust HTR in the rats (Fig. 2A-B). The HTR time-courses observed for the four higher LPH-5 doses provided interesting information about its pharmacokinetic properties (Fig. 2A). The rapid on-set of HTR following the LPH-5 injections indicates that the drug rapidly reaches the brain, with the 3.0, 6.0 and 12.0 mg/kg doses producing substantially higher HTR counts in the first 10-min interval after injection (0-10 min) than the 1.5 mg/kg dose. This dose-dependency was also reflected in the time profiles, as the HTR/10-min counts produced by 1.5 mg/kg LPH-5 peaked in the “20-30 min” interval, reflecting a CNS absorption component for this lower LPH-5 dose. In contrast, HTR/10-min counts for the intermediate 3.0 and 6.0 mg/kg doses peaked immediately after injection (“0-10 min”) and decreased incrementally from these peaks throughout the 180 min period. The dose differences were manifested in the different peak sizes and in the parallel-shifted time courses, which meant that HTR in rats dosed with 1.5 mg/kg LPH-5 leveled out after 120 min, whereas the two intermediate doses also produced substantial HTR/10-min counts in the 120-180 min interval. In animals dosed with 12.0 mg/kg LPH-5 rapidly established robust HTR levels that plateaued for essentially 120 min were observed, after which HTR/10-min counts incrementally decreased during the last 60 min (Fig. 2A). It is tempting to ascribe this profile for this highest drug dose to a compartment/bolus effect, which also is visible from the cumulative HTR time course (Fig. 2B). Interestingly, rats dosed with DOI (3 mg/kg) showed a markedly slower onset of HTR than the LPH-5-dosed animals, with the peak HTR plateau being in the “40-90” min interval (Fig. 2A).

Considering the different HTR time-course profiles observed for different LPH-5 doses, the dose-response relationship of the drug will inevitably be dependent on the time interval used for the analysis. Because of the longer lasting HTR induced by higher doses of the drug, the fitted ED_50_ values for LPH-5 based on the “0-30 min”, “0-60 min”, “0-120 min” and “0-180 min” data were 1.5 mg/kg, 1.3 mg/kg, 1.8 mg/kg and 3.0 mg/kg, respectively. Interestingly, this time- dependency also meant that DOI with its slower HTR induction displayed a lower HTR induction compared to that produced by 1.5 mg/kg LPH-5 based on “0-30 min” data, but a 2-fold higher HTR count than that of 1.5 mg/kg LPH-5 based on “0-180 min” data (Fig. 2C). It is not surprising that the peak responses produced by low and high doses of a drug will manifest themselves at different time points after drug administration, nor that that the relative efficacies exhibited by drugs with different response time-courses will depend on the recording time. We propose that this time-dependency is an important general aspect to consider when assessing and comparing dose-response relationships of drugs in their HTR induction.

### LPH-5 displays acute antidepressant-like effects in two rat models

The acute *in vivo* effects of LPH-5 administration were studied in two rat models of depression, using the 1.5 mg/kg (*i.p.*) dose of the drug found to induce robust but sub-maximal HTR levels in the SD rats (Fig. 2).

The FSL rat is a genetic rodent model of depression, as this animal in contrast to the control FRL rat exhibits depressive-like symptoms such as reduced appetite, reduced mobility, anhedonia, and sleep abnormalities.^43,44^ The potential locomotor activity effects of LPH-5 (1.5 mg/kg, *i.p.*) and the rapid-acting antidepressant drug *S-*ketamine (15 mg/kg, *i.p.*) were studied in an OFT, where the LPH-5-dosed rats did not display significantly changed locomotor activity compared to FSL rats, whereas both the FRL rats and the FSL rats treated with *S-*ketamine displayed decreased locomotor activity compared to the FSL animals [F(3,34)=3.990, p=0.0085 and p=0.0333, respectively] (Fig. 3A). In agreement with previous reports,^44,45^ the FSL rats exhibited significantly lower shorter periods of climbing [F(3,34)=28.69, p<0.0001; Dunnett’s p<0.0001] and significant longer periods of immobility [F(3,34)=27.88, p<0.0001; Dunnett’s p<0.0001] compared to the FRL animals in the FST, whereas time spent swimming did not differ significantly between animals in the two groups [F(3,34)=7.734, p=0.0005; Dunnett’s p=0.9627] (Fig. 3B). The immobility of the FSL animals was almost completely restored 1 h after injection of *S*-ketamine (p<0.0001), and LPH-5 also significantly reduced the immobility of the FSL animals (p=0.0194) (Fig. 3B). Only *S*-ketamine, not LPH-5, mediated a significant effect on the swimming (p=0.0003) and climbing (p=0.0422) times. Importantly, the reduced locomotor activity in the OFT combined with the increased mobility in the FST observed for *S*-ketamine is not seen as a systematic issue since the two effects of the drug act in opposite directions.

**Figure 3.**
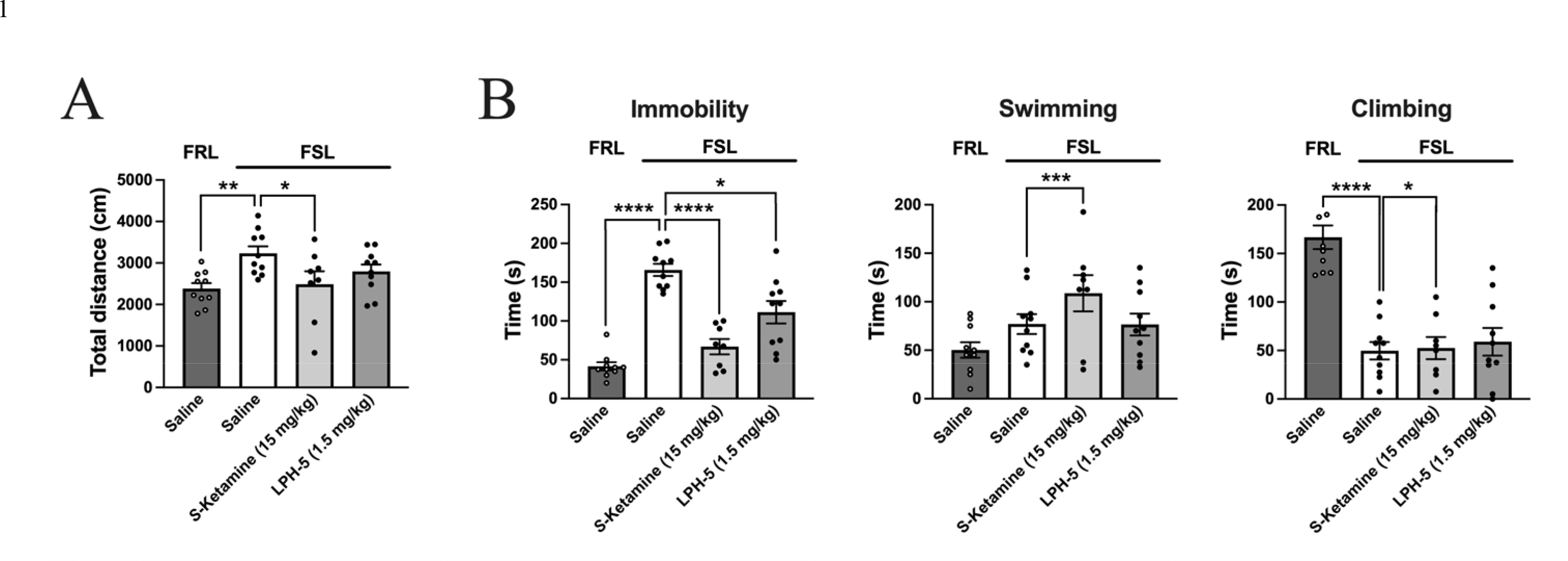
Acute antidepressant-like effects induced by LPH-5 in the Flinders Sensitive Line rat model of depression. The effects of *S*-ketamine and LPH-5 were investigated in the Open Field Test (OFT) and the Forced Swim Test (FST) performed 50 min and 60 min after the injections, respectively. **(A)** Effects of *i.p.* injections of saline, *S*-ketamine (15 mg/kg) and LPH-5 (1.5 mg/kg) in male Flinders Sensitive Line (FSL) rats on immobility, swimming and climbing (in a 5 min-period) in the FST. **(B)** Effects of *i.p.* injections of saline, *S*-ketamine (15 mg/kg) and LPH-5 (1.5 mg/kg) in male FSL rats on the total distance travelled (in a 5 min-period) in the OFT. (A-B) Male Flinders Resistant Line (FRL) rats were included as control rats. (A-B) Data are given as mean ± S.E.M. n=10 pr group, except for the *S*-ketamine group (n=8).

The putative antidepressant-like effects of LPH-5 (1.5 mg/kg, *i.p.*) were also studied in the ACTH-model, in which the disordered signaling in the hypothalamic–pituitary–adrenal (HPA) axis caused by repeated ACTH-dosing of the animals previously has been proposed to induce a depression-like state not treatable with classical antidepressants.^39,40^ The ACTH-mediated induction of this treatment-resistant depression was initially confirmed by use of the tricyclic antidepressant imipramine. In the FST, imipramine (15 mg/kg, *i.p.*) induced significant effects on the immobility of the saline-treated animals (p<0.0001, post-hoc analysis, two-way ANOVA), but not of the ACTH-treated animals (p=0.2060, post-hoc analysis, two-way ANOVA) (Fig. 4A *left*). In contrast, imipramine reduced locomotor activity in both saline- and ACTH-treated animals (p<0.0001 in both, post-hoc analysis, two-way ANOVA) (Fig. 4A *right*). Next, the effects of psilocybin (1 and 10 mg/kg, *i.p.*) and LPH-5 (1.5 mg/kg, *i.p.*) on ACTH-treated animals were studied (Fig. 4B and 4C). In the OFT, both psilocybin doses reduced the locomotor activity significantly (p=0.0194, and p<0.0001, post-hoc analysis, one-way ANOVA, respectively), whereas LPH-5 (1.5 mg/kg, *i.p.*) did not affect this parameter significantly (Fig 4B). In the FST, psilocybin (1 and 10 mg/kg, *i.p.*) did not significantly affect the immobility, swimming, or climbing times in the rats, whereas animals administered LPH-5 (1.5 mg/kg, *i.p.*) displayed a significant reduction in the immobility time (p=0.0183, post-hoc analysis, one-way ANOVA) but not in the swimming or climbing times compared to the saline-injected animals (Fig. 4C). It is a distinct possibility that different impact of psilocybin and LPH-5 on locomotor activity in the rats (Fig. 4B) contribute to their differential effects on the immobility time of the animals in the FST.

**Figure 4.**
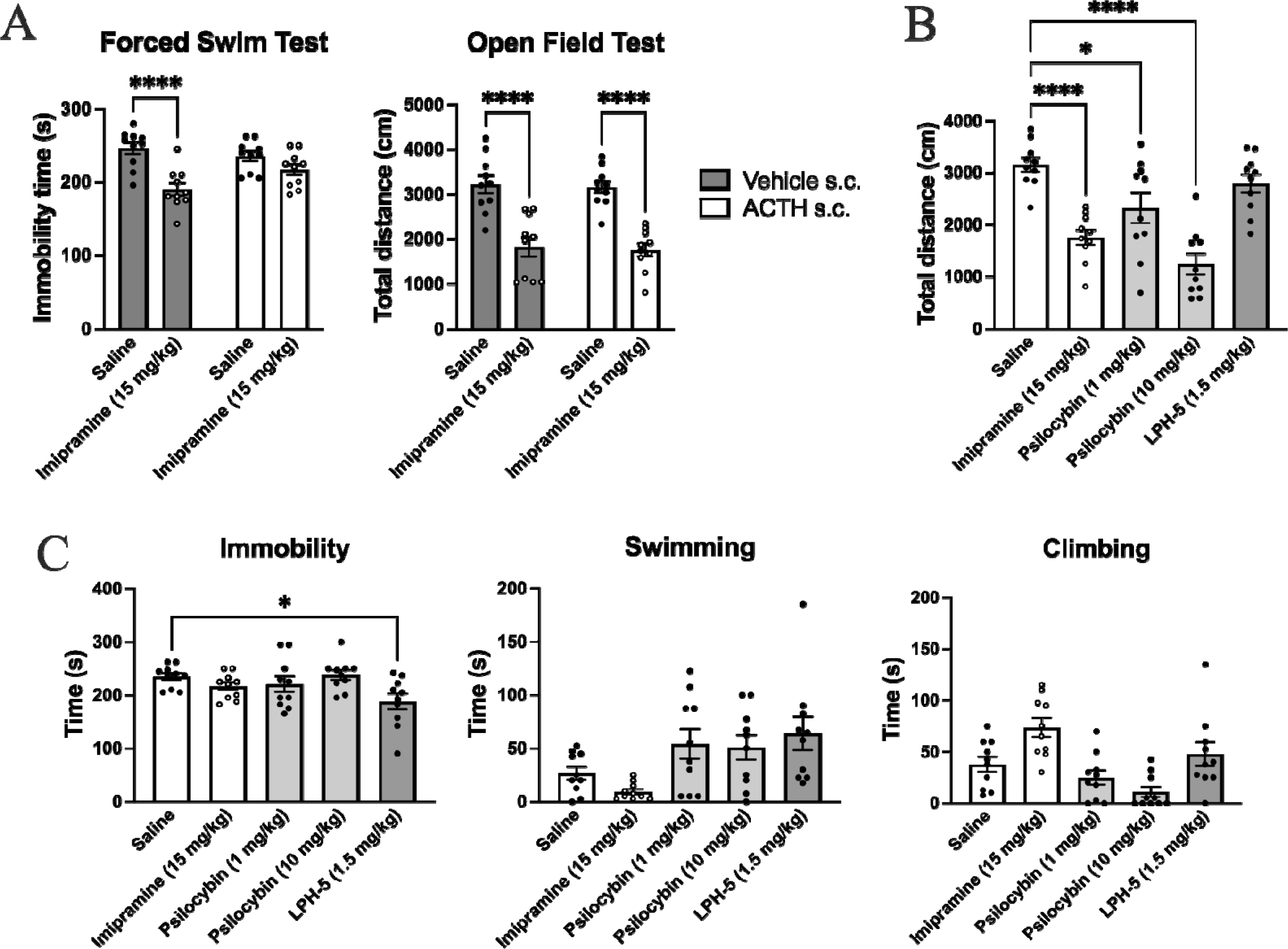
Acute antidepressant-like effects induced by LPH-5 in the ATCH rat model of depression. The effects of imipramine, psilocybin and LPH-5 in male Sprague Dawley rats were investigated in the Open Field Test (OFT) and the Forced Swim Test (FST) performed 50 min and 60 min after the injections, respectively. **(A)** Effects of *i.p.* injections of imipramine (15 mg/kg) in saline- and ACTH-treated rats on the immobility (in a 5 min-period) in the FST and on distance travelled (in a 5 min-period) in the OFT. Data are given as mean ± S.E.M. and are based on ten animals in each group (n=10). **(B)** Effects of *i.p.* injections of saline, imipramine (15 mg/kg), psilocybin (1.0 and 10 mg/kg) and LPH-5 (1.5 mg/kg) in ACTH-treated rats on the climbing, swimming, and immobility (in a 5 min-period) in the FST. Data are given as mean ± S.E.M. and are based on ten animals in each group (n=10). **(C)** Effects of *i.p.* injections of saline, imipramine (15 mg/kg), psilocybin (1.0 and 10 mg/kg) and LPH-5 (1.5 mg/kg) in ACTH-treated rats on the total distance travelled (in a 5 min-period) in the OFT. (A-C) Data are given as mean ± S.E.M. and are based on ten animals in each group (n=10).

### LPH-5 displays sustained antidepressant-like effects in a rat model

The effects of LPH-5 (0.3 and 1.5 mg/kg, *i.p.*) were tested in a previously developed model testing antidepressant-like effects of drugs in Wistar Kyoto rats 35 days after a single administration. Based on the antidepressant-like effects produced by psilocybin and LSD in this model, it has been proposed to reflect the long-lasting antidepressant effects of classical psychedelics in humans.^46^

In concordance with these previous findings,^46^ rats treated with psilocybin (1 mg/kg, *i.p.*) displayed significantly reduced immobility [F(3,28)=5.090, p=0.0061; post hoc p=0.0231 and significantly increased time spent swimming [F(3,28)=8.246, p=0.0004; post hoc p=0.0019] compared to the saline-treated rats (Fig. 5A). Analogously, both doses of LPH-5 (0.3 and 1.5 mg/kg, *i.p.*) also mediated significant reductions in the immobility of the rats [F(3,28)=5.090, p=0.0061; post hoc (0.3) p=0.0231 and (1.5) p=0.0117] and significant increases in the periods of swimming [F(3,28)=8.246, p=0.0004; post hoc (0.3) p=0.0019 and (1.5) p=0.0019], whereas neither psilocybin nor LPH-5 induced significant changes in the climbing parameter in the FST (Fig. 5A). None of the two drugs induced significant effects in the OFT, which suggests that the observations for them in the FST are not confounded by increased locomotor activity in the rats (Fig. 5B).

**Figure 5.**
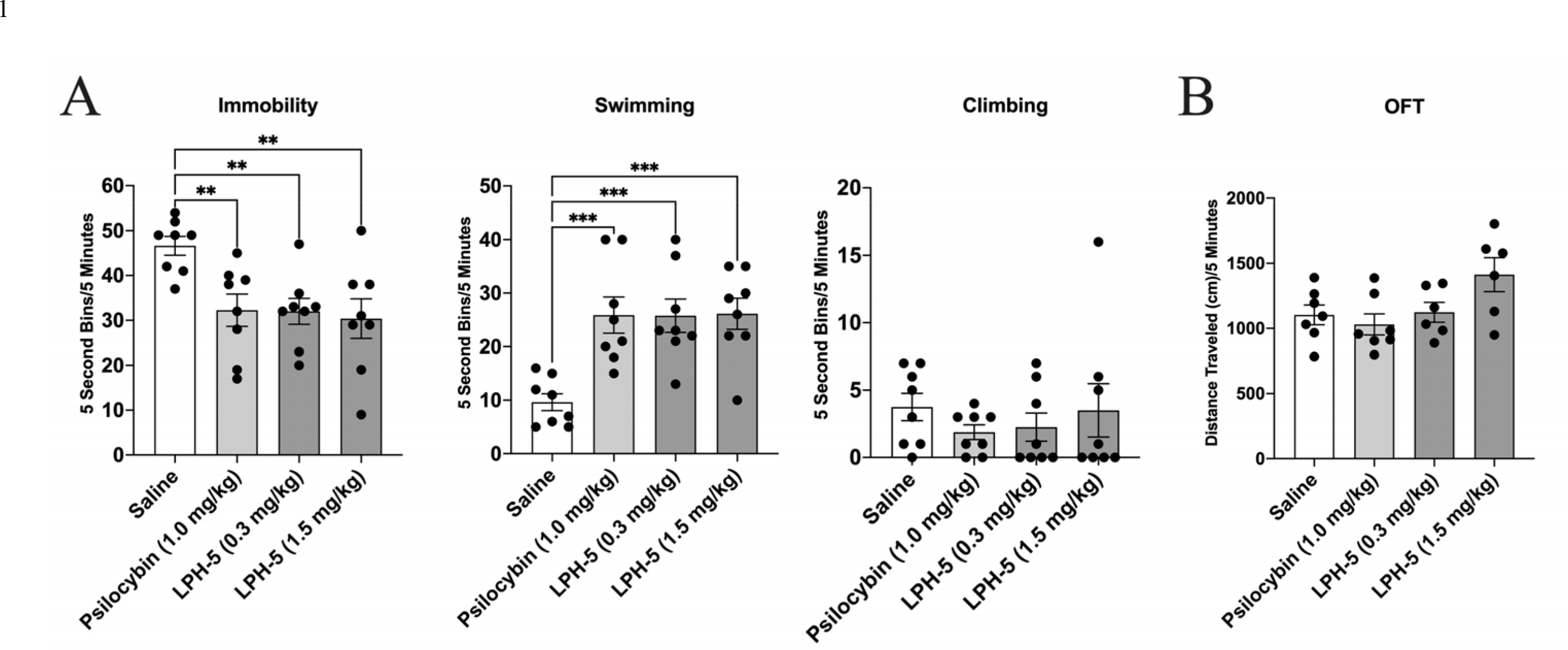
Long-term antidepressant-like effects induced by LPH-5 in rats. Forced Swim Test (FST) and locomotor activity studies were performed 35 days after a single *i.p.* injection of saline, psilocybin (1 mg/kg) or LPH-5 (0.3 and 1.5 mg/kg) in male Wistar Kyoto rats. **(A)** Effects of saline, psilocybin (1 mg/kg) and LPH-5 (0.3 and 1.5 mg/kg) on the immobility, swimming and climbing (5 sec bins in a 5 min-period) behaviors in the FST. **(B)** Effects of saline, psilocybin (1 mg/kg) and LPH-5 (0.3 and 1.5 mg/kg) on locomotor activity. Data are given as mean ± S.E.M. values and are based on 8 animals in each group (A) or 6-7 animals in each group (B). (**p<0.01; ***p<0.001; one way ANOVA with Bonferroni post-hoc test for multiple comparisons).

## DISCUSSION

In the present work we report that LPH-5 is one of a few truly selective 5-HT_2A_R agonists published to date, and taking advantage of its pharmacological profile we demonstrate that selective 5-HT_2A_R activation induces robust antidepressant-like effects in three rat models.

LPH-5 is a potent partial 5-HT_2A_R agonist displaying low-nanomolar EC_50_ values and R_max_ values of 56-94 % of the 5-HT R_max_ at the receptor in a battery of assays. Moreover, LPH-5 exhibits substantial selectivity towards 5-HT_2B_R and 5-HT_2C_R with its 10-60- and 10-100-fold lower binding affinities and functional potencies at 5-HT_2B_R and 5-HT_2C_R, respectively, than at 5-HT_2A_R (Fig. 1, Table 1). In contrast to its high agonist efficacy at 5-HT_2A_R and 5-HT_2B_R, its intrinsic activity at 5-HT_2C_R varies considerably in the five functional assays, spanning from robust agonist efficacy (inositol phosphate) over very low degrees of agonist efficacy (GTPγS binding and β-arrestin recruitment) to insignificant agonist efficacy (Ca^2+^ imaging and internalization) (Fig. 1, Table 1). Importantly, LPH-5 acts as a *de facto* antagonist of the 5-HT-induced response through 5-HT_2C_R in all four assays where it exhibits low intrinsic agonist activities at the receptor (Fig. 1, Table 1). This strongly suggests that LPH-5 is a very low- efficacious 5-HT_2C_R agonist, but that its efficacy is dependent on the nature and the sensitivity of the specific assay. Thus, LPH-5 both displays potency-based and functional selectivity within the 5-HT_2_Rs, and we propose that it potentially could exert inhibition rather than activation of 5- HT_2C_R *in vivo* when present in sufficiently high concentrations to target this subtype. All in all, the detailed pharmacological characterization of LPH-5 at recombinant receptors presented here and in our recent study^28^ collectively strongly supports its overall 5-HT_2A_R selectivity.

Judging from the comparable agonist potencies and efficacies displayed by LPH-5 at 5-HT_2A_R in assays for Gq- and β-arrestin-mediated signaling, the compound is a balanced agonist of the receptor (Fig. 1, Table 1). It has yet to be unequivocally addressed whether the antidepressant and other beneficial effects and the psychedelic effects mediated by the serotonergic psychedelics can be assigned to distinct 5-HT_2A_R signaling pathways, or whether the clinical efficacy of the drugs arise from and are in fact dependent on a non-biased activation of the receptor.

LPH-5 administration of LPH-5 in rats produced robust HTR, and both HTR counts and the time course of these over the recording period were dose-dependent, with a significant bolus effect observed for higher drug doses (Fig. 2). While it seems reasonable to consider the LPH-5- induced HTR a behavioral read-out of its 5-HT_2A_R activation, the HTR counts recorded at a specific time point does not directly correlate with its CNS exposure or its 5-HT_2A_R occupancy at that time, since there most likely is a temporal shift between agonist-mediated 5-HT_2A_R activation and the emergence of HTR. Nevertheless, we propose that the HTR data represents a reasonable approximation of the time course of *in vivo* 5-HT_2A_R engagement. The study of drug- induced HTR induction presented here is one of the most elaborate performed to date when it comes to the recording time, as previous studies of drug-induced HTR typically have employed recording times of 20-60 min.^47–49^ We propose that the detailed elucidation of the dose- dependency and the time course of the LPH-5-mediated HTR induction provide important information about its pharmacokinetic properties. The negligible and robust HTR induction mediated by 0.375 mg/kg and 1.5 mg/kg LPH-5, respectively, and its pharmacological profile at recombinant 5-HT_2_Rs and other serotonergic/monoaminergic receptors are of specific importance for this study, because these observations collectively indicate that the 1.5 mg/kg dose of the drug used in the behavioral studies will give rise to robust and selective 5-HT_2A_R activation.

The acute antidepressant-like effects produced by LPH-5 (1.5 mg/kg, *i.p.*) in the two rat models in this study align with the HTR data and further support that administration of this LPH-5 dose yield rapid and substantial CNS exposure levels of the drug (Figs. 3 and 4). Considering the absence of an effect of LPH-5 (1.5 mg/kg, *i.p.*) on locomotor activity of the rats, it is notable that this LPH-5 dose partially reverses the depression-like phenotype of FSL rats and induces a significant antidepressant-like effect in ACTH-treated rats (Figs. 3 and 4). While we will refrain from speculations as to which extent these acute antidepressant-like effects induced by LPH-5 in these models will translate into antidepressant efficacy in humans, it is notable that the drug thus exhibits antidepressant-like effects in both the genetic and the chemical-induced depression model.

Analogously to LSD and psilocybin, LPH-5 mediated robust antidepressant-like effects in a rat model proposed to reflect the long-lasting antidepressant effects produced by psychedelics in humans (Fig. 5).^46^ In view of the negligible HTR induction produced by LPH-5 (0.375 mg/kg, *i.p.*) in rats, it is notable that LPH-5 (0.3 mg/kg, *i.p.*) induces an antidepressant-like effect comparable to that produced by the higher dose (1.5 mg/kg, *i.p.*) (Figs. 2 and 5). This raises the question as to which extent the dose-HTR relationship exhibited by LPH-5 mirrors the *in vivo* receptor occupancy of the drug and whether activation of a low fraction of 5-HT_2A_Rs too low to induce substantial HTR could produce other effects *in vivo*. The effects of psychedelic microdosing are currently under intensive investigation,^50–53^ and substantial evidence certainly support that psychedelics even at sub-hallucinogenic doses can promote neurogenesis and neuroplasticity.^13,19,53–55^ The significant efficacy of a LPH-5 dose that does not elicit HTR in this rat depression model is intriguing, since it suggests that it may be possible to separate the putative long-term antidepressant-like effects from the acute psychedelic effects of the drug at low concentrations/doses. However, this speculation is largely based on assumed correlations between the LPH-5-induced HTRs and antidepressant-like effects in rats and the putative psychedelic and antidepressant effects of the drug in humans, respectively, that do not necessarily apply. Thus, the true test of whether the putative psychedelic and antidepressant effects of LPH-5 in humans can be separated by dose will be the clinical testing of the drug.

In conclusion, LPH-5 is a potent and selective 5-HT_2A_R agonist that induces HTR in a dose- dependent manner and mediates robust acute and persistent antidepressant-like effects in rodent models following a single administration. These findings corroborate the therapeutic potential of not only classical psychedelics characterized by high degrees of polypharmacology, but also selective 5-HT_2A_R agonists. We propose that LPH-5 could be a valuable pharmacological tool for investigations into the *in vivo* effects produced by selective 5-HT_2A_R activation and potentially a future therapeutic tool. At present, LPH-5 has been qualified for first-in-human trials, and we are poised to investigate if the promising results for the drug presented here will translate into clinical efficacy.

## Acknowledgments

The FRL/FSL rats were donated by Dr. Gregers Wegener (Translational Neuropsychiatry Unit, Aarhus University).

## Funding

Generous support from the Lundbeck Foundation (R208-2015-3140), Innovations Fund Denmark (0174-00076A) and Novo Nordisk Foundation (NNF19OC 0058080) is gratefully acknowledged.

### Author contributions

Conceptualization: AAJ, EMR, CDN, BE, JLK Methodology: CRC, MH, AHB, CDN, BE Investigation: AAJ, CRC, MH, AHB, EK, BE, CDN Visualization: AAJ, CDN, BE

Project administration: AAJ, EMR, JLK Supervision: BE, CDN

Writing – original draft: AAJ, CDN, BE, JLK

Writing – review & editing: AAJ, CRC, MH, AHB, EK, EMR, CDN, BE, JLK

## Supplementary Information

Supplementary information is available at MP’s website.

## References

1. Acero VP, Cribas ES, Browne KD, Rivellini O, Burrell JC, O’Donnell JC et al. Bedside to bench: the outlook for psychedelic research. Front Pharmacol 2023; 14: 1240295.

2. Carhart-Harris RL, Bolstridge M, Day CMJ, Rucker J, Watts R, Erritzøe DE et al. Psilocybin with psychological support for treatment-resistant depression: six-month follow-up. Psychopharmacology (Berl) 2018; 235(2): 399–408.

3. Reiff CM, Richman EE, Nemeroff CB, Carpenter LL, Widge AS, Rodriguez CI et al. Psychedelics and Psychedelic-Assisted Psychotherapy. Am J Psychiatry 2020; 177(5): 391–410.

4. Gukasyan N, Schreyer CC, Griffiths RR, Guarda AS. Psychedelic-Assisted Therapy for People with Eating Disorders. Curr Psychiatry Rep 2022; 24(12): 767–775.

5. Yaden DB, Nayak SM, Gukasyan N, Anderson BT, Griffiths RR. The Potential of Psychedelics for End of Life and Palliative Care. Curr Top Behav Neurosci 2022; 56: 169–184.

6. Andersen KAA, Carhart-Harris R, Nutt DJ, Erritzøe D. Therapeutic effects of classic serotonergic psychedelics: A systematic review of modern-era clinical studies. Acta Psychiatr Scand 2021; 143(2): 101–118.

7. Griffiths RR, Johnson MW, Carducci MA, Umbricht A, Richards WA, Richards BD et al. Psilocybin produces substantial and sustained decreases in depression and anxiety in patients with life-threatening cancer: A randomized double-blind trial. J Psychopharmacol 2016; 30(12): 1181–1197.

8. McClure-Begley TD, Roth BL. The promises and perils of psychedelic pharmacology for psychiatry. Nat Rev Drug Discov 2022; 21(6): 463–473.

9. Nichols DE. Psychedelics. Pharmacol Rev 2016; 68(2): 264–355.

10. Moliner R, Girych M, Brunello CA, Kovaleva V, Biojone C, Enkavi G et al. Psychedelics promote plasticity by directly binding to BDNF receptor TrkB. Nat Neurosci 2023; 26(6): 1032–1041.

11. Gumpper RH, Roth BL. Psychedelics: preclinical insights provide directions for future research. Neuropsychopharmacology 2023.

12. Jaster AM, González-Maeso J. Mechanisms and molecular targets surrounding the potential therapeutic effects of psychedelics. Mol Psychiatry 2023.

13. Kwan AC, Olson DE, Preller KH, Roth BL. The neural basis of psychedelic action. Nat Neurosci 2022; 25(11): 1407–1419.

14. Carhart-Harris RL, Nutt DJ. Serotonin and brain function: a tale of two receptors. J Psychopharmacol 2017; 31(9): 1091–1120.

15. Inserra A, Piot A, De Gregorio D, Gobbi G. Lysergic Acid Diethylamide (LSD) for the Treatment of Anxiety Disorders: Preclinical and Clinical Evidence. CNS Drugs 2023; 37(9): 733–754.

16. Cameron LP, Benetatos J, Lewis V, Bonniwell EM, Jaster AM, Moliner R et al. Beyond the 5-HT_2A_ Receptor: Classic and Nonclassic Targets in Psychedelic Drug Action. J Neurosci 2023; 43(45): 7472–7482.

17. Wallach J, Cao AB, Calkins MM, Heim AJ, Lanham JK, Bonniwell EM et al. Identification of 5-HT_2A_ Receptor Signaling Pathways Responsible for Psychedelic Potential. bioRxiv 2023.

18. Cao D, Yu J, Wang H, Luo Z, Liu X, He L et al. Structure-based discovery of nonhallucinogenic psychedelic analogs. Science 2022; 375(6579): 403-411.

19. Vargas MV, Dunlap LE, Dong C, Carter SJ, Tombari RJ, Jami SA et al. Psychedelics promote neuroplasticity through the activation of intracellular 5-HT_2A_ receptors. Science 2023; 379(6633): 700-706.

20. Zhang G, Stackman RW, Jr. The role of serotonin 5-HT_2A_ receptors in memory and cognition. Front Pharmacol 2015; 6: 225.

21. Meltzer HY, Massey BW, Horiguchi M. Serotonin receptors as targets for drugs useful to treat psychosis and cognitive impairment in schizophrenia. Curr Pharm Biotechnol 2012; 13(8): 1572–1586.

22. Jalal B. The neuropharmacology of sleep paralysis hallucinations: serotonin 2A activation and a novel therapeutic drug. Psychopharmacology (Berl) 2018; 235(11): 3083–3091.

23. Simon IA, Bjørn-Yoshimoto WE, Harpsøe K, Iliadis S, Svensson B, Jensen AA et al. Ligand selectivity hotspots in serotonin GPCRs. Trends Pharmacol Sci 2023.

24. Hansen M, Phonekeo K, Paine JS, Leth-Petersen S, Begtrup M, Bräuner-Osborne H et al. Synthesis and structure-activity relationships of N-benzyl phenethylamines as 5-HT_2A/2C_ agonists. ACS Chem Neurosci 2014; 5(3): 243–249.

25. Jensen AA, McCorvy JD, Leth-Petersen S, Bundgaard C, Liebscher G, Kenakin TP et al. Detailed Characterization of the In Vitro Pharmacological and Pharmacokinetic Properties of N-(2-Hydroxybenzyl)-2,5-Dimethoxy-4-Cyanophenylethylamine (25CN-NBOH), a Highly Selective and Brain-Penetrant 5-HT_2A_ Receptor Agonist. J Pharmacol Exp Ther 2017; 361(3): 441-453.

26. Kim K, Che T, Panova O, DiBerto JF, Lyu J, Krumm BE et al. Structure of a Hallucinogen-Activated Gq-Coupled 5-HT_2A_ Serotonin Receptor. Cell 2020; 182(6): 1574–1588.e1519.

27. Kaplan AL, Confair DN, Kim K, Barros-Álvarez X, Rodriguiz RM, Yang Y et al. Bespoke library docking for 5-HT_2A_ receptor agonists with antidepressant activity. Nature 2022; 610(7932): 582-591.

28. Rørsted EM, Jensen AA, Smits G, Frydenvang KA, Kristensen JL. Discovery and Structure-Activity Relationships of 2,5-dimethoxyphenylpiperidines as Serotonin 5-HT_2A_ Receptor Agonists. J Med Chem. Accepted.

29. Kristensen JL, Jensen AA, Märcher-Rørsted E, Leth-Petersen S. 5-HT2A agonists for use in the treatment of depression. vol. US 11,246,860 B22022.

30. Trinquet E, Fink M, Bazin H, Grillet F, Maurin F, Bourrier E et al. D-myo-inositol 1- phosphate as a surrogate of D-myo-inositol 1,4,5-tris phosphate to monitor G protein-coupled receptor activation. Anal Biochem 2006; **358**(1): 126-135.

31. Jensen AA, Plath N, Pedersen MH, Isberg V, Krall J, Wellendorph P et al. Design, synthesis, and pharmacological characterization of N- and O-substituted 5,6,7,8- tetrahydro-4H-isoxazolo[4,5-d]azepin-3-ol analogues: novel 5-HT_2A_/5-HT_2C_ receptor agonists with pro-cognitive properties. J Med Chem 2013; **56**(3): 1211-1227.

32. Golla R, Seethala R. A homogeneous enzyme fragment complementation cyclic AMP screen for GPCR agonists. J Biomol Screen 2002; 7(6): 515–525.

33. Wang T, Li Z, Cvijic ME, Zhang L, Sum CS. Measurement of cAMP for G(αs)- and G(αi) Protein-Coupled Receptors (GPCRs). In: Markossian S, Grossman A, Brimacombe K, Arkin M, Auld D, Austin C et al. (eds). Assay Guidance Manual. Eli Lilly & Company and the National Center for Advancing Translational Sciences: Bethesda (MD), 2004.

34. Johnson EN, Shi X, Cassaday J, Ferrer M, Strulovici B, Kunapuli P. A 1,536-well [^35^S]GTPgammaS scintillation proximity binding assay for ultra-high-throughput screening of an orphan galphai-coupled GPCR. Assay Drug Dev Technol 2008; 6(3): 327–337.

35. Muneta-Arrate I, Diez-Alarcia R. [^35^S]GTPγS (Guanosine-5’-O-(γ-thio)triphosphate- [^35^S]) Binding Scintillation Proximity Assay Experiments in Postmortem Brain Tissue. Methods Mol Biol 2023; 2687: 31–43.

36. Bouma J, Soethoudt M, van Gils N, Xia L, van der Stelt M, Heitman LH. Cellular Assay to Study β-Arrestin Recruitment by the Cannabinoid Receptors 1 and 2. Methods Mol Biol 2023; 2576: 189–199.

37. Nakane A, Gotoh Y, Ichihara J, Nagata H. New screening strategy and analysis for identification of allosteric modulators for glucagon-like peptide-1 receptor using GLP-1 (9-36) amide. Anal Biochem 2015; 491: 23–30.

38. Detke MJ, Rickels M, Lucki I. Active behaviors in the rat forced swimming test differentially produced by serotonergic and noradrenergic antidepressants. Psychopharmacology (Berl) 1995; 121(1): 66–72.

39. Pereira VS, Joca SRL, Harvey BH, Elfving B, Wegener G. Esketamine and rapastinel, but not imipramine, have antidepressant-like effect in a treatment-resistant animal model of depression. Acta Neuropsychiatr 2019; 31(5): 258–265.

40. Kaadt E, Højgaard K, Mumm B, Christiansen SL, Müller HK, Damgaard CK et al. Dysregulation of miR-185, miR-193a, and miR-450a in the skin are linked to the depressive phenotype. Prog Neuropsychopharmacol Biol Psychiatry 2021; 104: 110052.

41. Halberstadt AL, Geyer MA. Effect of Hallucinogens on Unconditioned Behavior. Curr Top Behav Neurosci 2018; 36: 159–199.

42. Odland AU, Kristensen JL, Andreasen JT. Animal Behavior in Psychedelic Research. Pharmacol Rev 2022; 74(4): 1176–1205.

43. Overstreet DH, Friedman E, Mathé AA, Yadid G. The Flinders Sensitive Line rat: a selectively bred putative animal model of depression. Neurosci Biobehav Rev 2005; 29(4- 5): 739–759.

44. Overstreet DH, Wegener G. The flinders sensitive line rat model of depression--25 years and still producing. Pharmacol Rev 2013; 65(1): 143–155.

45. du Jardin KG, Liebenberg N, Cajina M, Müller HK, Elfving B, Sanchez C et al. S-Ketamine Mediates Its Acute and Sustained Antidepressant-Like Activity through a 5- HT_1B_ Receptor Dependent Mechanism in a Genetic Rat Model of Depression. Front Pharmacol 2017; 8: 978.

46. Hibicke M, Landry AN, Kramer HM, Talman ZK, Nichols CD. Psychedelics, but Not Ketamine, Produce Persistent Antidepressant-like Effects in a Rodent Experimental System for the Study of Depression. ACS Chem Neurosci 2020; 11(6): 864–871.

47. Abiero A, Botanas CJ, Sayson LV, Custodio RJ, de la Peña JB, Kim M et al. 5-Methoxy- α-methyltryptamine (5-MeO-AMT), a tryptamine derivative, induces head-twitch responses in mice through the activation of serotonin receptor 2a in the prefrontal cortex. Behav Brain Res 2019; 359: 828–835.

48. Klein AK, Chatha M, Laskowski LJ, Anderson EI, Brandt SD, Chapman SJ, et al. Investigation of the Structure-Activity Relationships of Psilocybin Analogues. ACS Pharmacol Transl Sci 2021; 4(2): 533-542.

49. Buchborn T, Lyons T, Knöpfel T. Tolerance and Tachyphylaxis to Head Twitches Induced by the 5-HT_2A_ Agonist 25CN-NBOH in Mice. Front Pharmacol 2018; 9: 17.

50. Wong A, Raz A. Microdosing with classical psychedelics: Research trajectories and practical considerations. Transcult Psychiatry 2022; 59(5): 675–690.

51. Kiilerich KF, Lorenz J, Scharff MB, Speth N, Brandt TG, Czurylo J et al. Repeated low doses of psilocybin increase resilience to stress, lower compulsive actions, and strengthen cortical connections to the paraventricular thalamic nucleus in rats. Mol Psychiatry 2023; 28(9): 3829–3841.

52. Glazer J, Murray CH, Nusslock R, Lee R, de Wit H. Low doses of lysergic acid diethylamide (LSD) increase reward-related brain activity. Neuropsychopharmacology 2023; 48(2): 418–426.

53. Calder AE, Hasler G. Towards an understanding of psychedelic-induced neuroplasticity. Neuropsychopharmacology 2023; **48**(1): 104-112.

54. de Vos CMH, Mason NL, Kuypers KPC. Psychedelics and Neuroplasticity: A Systematic Review Unraveling the Biological Underpinnings of Psychedelics. Front Psychiatry 2021; 12: 724606.

55. Ly C, Greb AC, Cameron LP, Wong JM, Barragan EV, Wilson PC et al. Psychedelics Promote Structural and Functional Neural Plasticity. Cell Rep 2018; 23(11): 3170–3182.

