## Supplementary material for "The selective 5-HT_2A_ receptor agonist LPH-5 induces persistent and robust antidepressant-like effects in rodents": SI

### Supplemental Information

- Detailed descriptions of the protocols for the functional assays
- Supplemental Figure 1

Text summaries and file types of the Supplemental Information:

*Detailed descriptions of the protocols for the functional assays.*

A detailed description of the assay protocols used. A Word file.

*Supplemental Figure 1.*

A figure over all HTR data (supplementary to Fig. 2 in the manuscript). Incorporated in a Word file (can also be provided in tif or jpg format).

**Detailed descriptions of the protocols for the functional assays**

***Inositol phosphate assay.*** The functional characterization of LPH-5 at human 5-$HT_{2A}$R, 5-$HT_{2B}$R and 5-$HT_{2C}$R in the IP-One HTRF® assay[1,2] (Cisbio, Bagnol, France) was performed by Eurofins. Briefly, HEK293 cells expressing the respective receptors were suspended in buffer (10 mM HEPES, 4.2 mM KCl, 146 mM NaCl, 1 mM $CaCl_2$, 0.5 mM $MgCl_2$, 5.5 mM glucose, 50 mM LiCl, pH 7.4) distributed in 384-well microplates at a density of 2 x $10^4$ cells/well and incubated for 30 min at 37°C in the presence of buffer (basal control) or the test compound. Following incubation, the cells were lysed and the fluorescence acceptor (D2-labeled $IP_1$) and fluorescence donor (anti-$IP_1$ antibody labeled with europium cryptate) were added. After 60 min at room temperature, the fluorescence transfer was measured at $\lambda$ex=337 nm and $\lambda$em=620 nm and 665 nm using an Envision™ microplate reader (Perkin Elmer, Boston, MA) The $IP_1$ concentration was determined by the ratio between the signal measured at 665 nm and that measured at 620 nm.

***cAMP assay.*** The functional characterization of LPH-5 at human 5-$HT_{1B}$R in a HitHunter® cAMP assay[3,4] was performed by Eurofins. Stable 5-$HT_{1B}$R-CHO-K1 cells were assayed in this Enzyme Fragment Complementation (EFC)-based assay, where a fragment β-galactosidase enzyme donor (ED) is conjugated with cAMP. This ED-cAMP conjugate and intracellular cAMP compete for binding to an anti-cAMP antibody (Ab), and thus the level of cAMP in the cell impacts the ability of the cAMP-ED conjugate to complement with fragment β-galactosidase acceptor (EA) and form the active enzyme that hydrolyzes a substrate to produce a chemiluminescent signal. Thus, the amount of cAMP in the cell is directly proportional with the signal. Briefly, stable 5-$HT_{1B}$R-CHO-K1 cells were seeded in a total volume of 20 µL into white walled, 384-well microplates and incubated at 37°C overnight. Prior to testing cell plating media was exchanged with 10 µL of assay buffer (HBSS + 10 mM HEPES, pH 7.4). 5 µL of compound solution and 5 µL of forskolin solution (assay concentration of forskolin: 15 µM) was added to cells and incubated for 30 min at 37°C. HitHunter ED-cAMP and anti-cAMP reagents were added followed by 1 h incubation at RT. Assay signal was generated through addition of HitHunter EA reagent and 2 h incubation at RT. The plates were assayed following signal generation in an Envision™ microplate reader (PerkinElmer) for chemiluminescent signal detection.

***GTPγS binding assays.*** The functional characterization of LPH-5 at human 5-$HT_{2A}$R and 5-$HT_{2C}$R in a Scintillation Proximity Assay (SPA)-based GTPγS binding assay[5,6] was performed by Eurofins. Briefly, membranes from stable 5-$HT_{2A}$R- and 5-$HT_{2C}$R-CHO-K1 cell lines were pre-incubated with 1 μM GDP and vehicle or test compound in assay buffer (5-$HT_{2A}$R: 20 mM HEPES, 30 mM NaCl, 10 mM $MgCl_2$, 1 mM DTT, 1 mM EDTA, 40 μg/ml saponin, pH 7.4; 5-$HT_{2C}$R: 20 mM HEPES, 100 mM NaCl, 10 mM $MgCl_2$, 1 mM DTT, 1 mM EDTA, pH 7.4) for 20 min, and then SPA beads were added for another 60 min incubation at 30 °C. The reaction was then initiated by addition of [$^{35}$S]GTPγS (0.3 nM) followed by 15 min incubation. After incubation, the binding of [$^{35}$S]GTPγS to G-protein was detected by MicroBeta Microplate Counter (PerkinElmer). For testing in agonist mode, the response induced by the test compound through the receptor was determined. For testing in antagonist mode at 5-$HT_{2C}$R, the inhibition mediated by the compound at the 5-HT (1 μM)-induced response through the receptor was determined.

***β-arrestin recruitment assays.*** The functional characterization of LPH-5 at human 5-$HT_{2A}$R, 5-$HT_{2C}$R and 5-$HT_{1B}$R in PathHunter® β-arrestin assays[7] was performed by Eurofins. Briefly, stable 5-$HT_{2A}$R-, 5-$HT_{2C}$R- and 5-$HT_{1B}$R-U2OS cell lines co-expressing ProLink™ (PK)-tagged receptor and Enzyme Acceptor (EA)-tagged β-arrestin were used, where receptor activation induces β-arrestin recruitment, thus forcing complementation of the two enzyme fragments (EA and PK), and the resulting functional enzyme hydrolyzes substrate to generate a chemiluminescent signal. Cells were seeded in a total volume of 20 μL into white-walled, 384-well microplates and incubated at 37 °C until testing. For testing in agonist mode at the three receptors, cells were incubated with sample containing the compound to induce response at 37°C for 120 min. For testing in antagonist mode at 5-$HT_{2C}$R, cells were pre-incubated with the compound for 30 min at 37°C, followed by agonist challenge (5-HT $EC_{80}$) and incubation for 120 min at 37°C. The assay signal was generated by addition of PathHunter Detection reagent cocktail followed by incubation for 1 h. The plates were assayed following signal generation in an Envision™ microplate reader (PerkinElmer) for chemiluminescent signal detection.

***Internalization assays.*** The functional characterization of LPH-5 at human 5-$HT_{2A}$R and 5-$HT_{2C}$R stably expressed in U2OS cells in PathHunter® internalization assays[8] was performed by Eurofins. The 5-$HT_{2A}$R was assayed in a PathHunter total GPCR internalization assay format using the 5-$HT_{2A}$R-U2OS cell line, where the ProLink™ (PK)-tagged receptor was co-expressed with an Enzyme Acceptor (EA)-tag localized to the endosomes. The 5-$HT_{2A}$R was assayed in a PathHunter activated GPCR internalization assay format using the 5-$HT_{2C}$R-U2OS cell line, where the receptor was co-expressed with EA-tagged β-arrestin, and a PK-tag localized to the endosomes. In both cell lines, receptor activation induces β-arrestin recruitment and internalization of the receptor/β-arrestin complex in the endosomes, which forces complementation of the two β-galactosidase enzyme fragments (EA and PK) thus forming a functional enzyme that hydrolyzes substrate to generate a chemiluminescent signal. For testing in agonist mode, cells were incubated with sample containing the compound to induce response for 180 min at 37°C. For testing in antagonist mode at 5-$HT_{2C}$R, cells were pre-incubated with the compound for 60 minutes at 37°C, followed by agonist challenge (5-HT $EC_{80}$) and incubation for 180 min at 37°C. The assay signal was generated by addition of PathHunter Detection reagent cocktail followed by incubation for 1 h. The plates were assayed following signal generation in an Envision™ microplate reader (PerkinElmer) for chemiluminescent signal detection.

***References***


1. Trinquet E, Fink M, Bazin H, Grillet F, Maurin F, Bourrier E *et al.* D-myo-inositol 1-phosphate as a surrogate of D-myo-inositol 1,4,5-tris phosphate to monitor G protein-coupled receptor activation. *Anal Biochem* 2006; **358**(1)**:** 126-135.

2. Jensen AA, Plath N, Pedersen MH, Isberg V, Krall J, Wellendorph P *et al.* Design, synthesis, and pharmacological characterization of N- and O-substituted 5,6,7,8-tetrahydro-4H-isoxazolo[4,5-d]azepin-3-ol analogues: novel 5-$HT_{2A}$/5-$HT_{2C}$ receptor agonists with pro-cognitive properties. *J Med Chem* 2013; **56**(3)**:** 1211-1227.

3. Golla R, Seethala R. A homogeneous enzyme fragment complementation cyclic AMP screen for GPCR agonists. *J Biomol Screen* 2002; **7**(6)**:** 515-525.

4. Wang T, Li Z, Cvijic ME, Zhang L, Sum CS. Measurement of cAMP for G(αs)- and G(αi) Protein-Coupled Receptors (GPCRs). In: Markossian S, Grossman A, Brimacombe K, Arkin M, Auld D, Austin C *et al.* (eds). *Assay Guidance Manual*. Eli Lilly & Company and the National Center for Advancing Translational Sciences: Bethesda (MD), 2004.

5. Johnson EN, Shi X, Cassaday J, Ferrer M, Strulovici B, Kunapuli P. A 1,536-well [(35)S]GTPgammaS scintillation proximity binding assay for ultra-high-throughput screening of an orphan galphai-coupled GPCR. *Assay Drug Dev Technol* 2008; **6**(3)**:** 327-337.

6. Muneta-Arrate I, Diez-Alarcia R. [(35)S]GTPγS (Guanosine-5'-O-(γ-thio)triphosphate-[(35)S]) Binding Scintillation Proximity Assay Experiments in Postmortem Brain Tissue. *Methods Mol Biol* 2023; **2687:** 31-43.

7. Bouma J, Soethoudt M, van Gils N, Xia L, van der Stelt M, Heitman LH. Cellular Assay to Study β-Arrestin Recruitment by the Cannabinoid Receptors 1 and 2. *Methods Mol Biol* 2023; **2576:** 189-199.

8. Nakane A, Gotoh Y, Ichihara J, Nagata H. New screening strategy and analysis for identification of allosteric modulators for glucagon-like peptide-1 receptor using GLP-1 (9-36) amide. *Anal Biochem* 2015; **491:** 23-30.

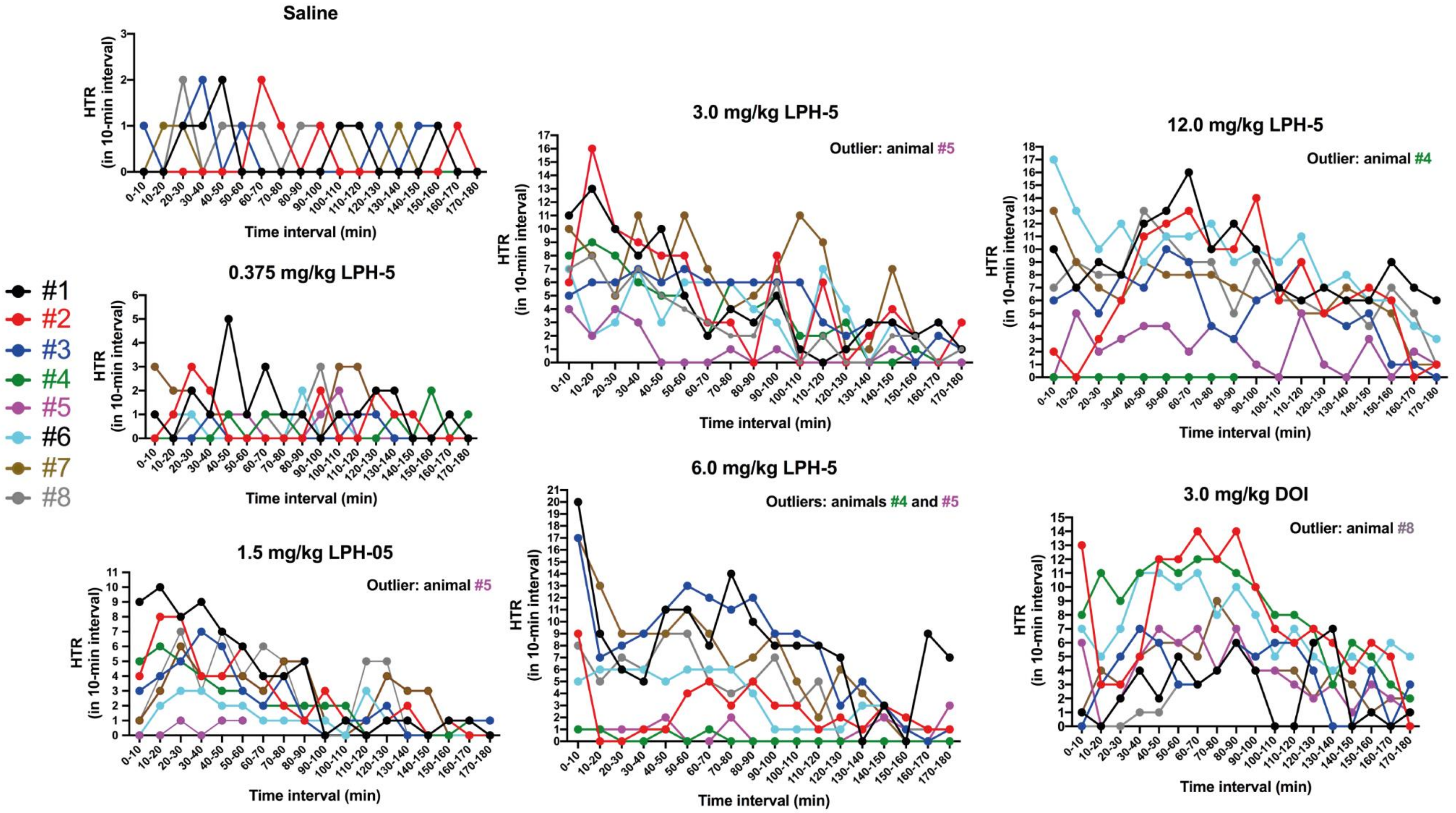


**Supplemental Figure 1.** Head twitch response (HTR) induced by *i.p.* injections of saline, LPH-05 (0.375; 1.5; 3.0; 6.0; 12.0 mg/kg) and DOI (3.0 mg/kg) in the individual Sprague Dawley rats, 8 animals per group. The specific HTR data registered for each individual rat for each 10-min interval over a 180 min-period following injection are given. The data from rats considered outliers not included in Fig. 2 are given in this figure, where the outlier animals are indicated.
